# The legacy of C_4_ evolution in the hydraulics of C_3_ and C_4_ grasses

**DOI:** 10.1101/2020.05.14.097030

**Authors:** Haoran Zhou, Erol Akçay, Erika Edwards, Brent Helliker

**Affiliations:** Department of Biology, University of Pennsylvania, Philadelphia, PA 19104, USA; Department of Ecology and Evolutionary Biology, Yale University, New Haven, CT 06511, USA; University Corporation for Atmospheric Research, Boulder, CO 80301, USA

**Author notes:** Correspondence*: Haoran Zhou, *Email. Erol Akçay, Erika Edwards, Brent R. Helliker.

## Abstract

The anatomical reorganization required for optimal C_4_ photosynthesis should also impact plant hydraulics. Most C_4_ plants possess large bundle-sheath cells and high vein density, which should also lead to higher leaf hydraulic conductance (K_leaf_) and capacitance. Paradoxically, the C_4_ pathway reduces water demand and increases water-use-efficiency, creating a potential mis-match between supply capacity and demand in C_4_ plant water relations. We use phylogenetic analyses, physiological measurements, and models to examine the reorganization of hydraulics in closely-related C_4_ and C_3_ grasses. Evolutionarily young C_4_ lineages have higher K_leaf_, capacitance, turgor-loss-point, and lower stomatal conductance than their C_3_ relatives. In contrast, species from older C_4_ lineages show decreased K_leaf_ and capacitance, indicating that over time, C_4_ plants have evolved to optimize hydraulic investments while maintaining C_4_ anatomical requirements. The initial “over-plumbing” of C_4_ plants disrupts the positive correlation between maximal assimilation rate and K_leaf_, decoupling a key relationship between hydraulics and photosynthesis generally observed in vascular plants.

## Introduction

The evolution of C_4_ photosynthesis in the grasses— and the attendant fine-tuning of both anatomical and biochemical components across changing selection landscapes^[1,2,3]^— likely impacted leaf hydraulics and hydraulics-photosynthesis relationships, both within the grass lineages in which C_4_ evolved independently > 20 times^[4]^, and as compared to closely-related C_3_^[5,6]^. C_4_ plants typically exhibit lower stomatal conductance (g_s_) and consequently greater water-use efficiency than C_3_, because the concentration of CO_2_ inside bundle sheath cells permits reduced intercellular CO_2_ concentrations and conservative stomatal behavior^[7,8,9]^. At the same time, C_4_ plants require high bundle sheath to mesophyll ratios (BS:M), which are accomplished with increased vein density and bundle sheath size as compared to C_3_ plants. In C_3_ species, leaf hydraulic conductance (K_leaf_) has a positive relationship with vein density^[10,11,12,13]^. The decreased inter-veinal distance and consequently higher vein density in C_4_ species has been predicted to lead to a higher K_leaf_ than closely-related C_3_ species^[14,15]^. Further, increased bundle sheath size was proposed to lead to a higher leaf capacitance in C_4_ species^[15,16]^, This would lead to a potential physiological “mis-match”, where the evolution of the C_4_ pathway simultaneously increases a plant’s hydraulic capacity while reducing its transpirational demand.

The significance of such a potential physiological mismatch depends on the potential costs and tradeoffs associated with the building of an ‘over-plumbed’ leaf. If the costs are high^[12,17]^, then one would expect to see a reduction of K_leaf_ over evolutionary time, as continued selection works to optimize the C_4_ metabolism^[5,18]^. Alternatively, a maintenance of high K_leaf_ over time could result from either a lack of strong selection to reduce K_leaf_, or a strong evolutionary constraint imposed by the anatomical requirements of C_4_ photosynthesis. In other words, the high BS:M ratio required for an efficient C_4_ system may directly limit the ability of C_4_ plants to optimize their hydraulic architecture.

The evolution of a new photosynthetic pathway that results in multiple potential changes to the plant hydraulic system represents the ideal platform to expand our understanding of the relationship between photosynthesis and water transport. It is generally thought that maximum photosynthetic rate (A_max_) and hydraulic capacity (K_leaf_) are tightly linked, because the ability to transport water through leaves to the sites of evaporation at a high rate allows for the maximization of carbon gain. Studies have documented a positive correlation between A_max_ and K_leaf_ across many scales, from a broad phylogenetic spectrum of species spanning vascular plants^[11]^, to smaller clades of closely related species^[13]^. Grasses are largely absent from previous efforts to examine this relationship, which is unfortunate because of the parallel venation found in grasses and other monocots. With over 20 origins of C_4_ photosynthesis with ages that span ~ 30 million years, grasses also present a unique opportunity to examine the influence of C_4_ evolution on A_max_-K_leaf_ relationships. Using a broad sampling of grasses (Fig. 1), we determined whether anatomical differences associated with C_4_ evolution result in greater K_leaf_ and leaf capacitance compared to their C_3_ relatives. We then compared these properties between closely related C_3_ and C_4_ clades to determine how C_4_ evolution alters the predicted A_max_-K_leaf_ relationships. Finally, we then quantified evolutionary trends in K_leaf_, capacitance and turgor loss point after the evolution of C_4_ within a lineage by asking whether more recent origins of C_4_ are represented by higher K_leaf_ and a greater K_leaf_-A_max_ mismatch.

**Fig. 1.**
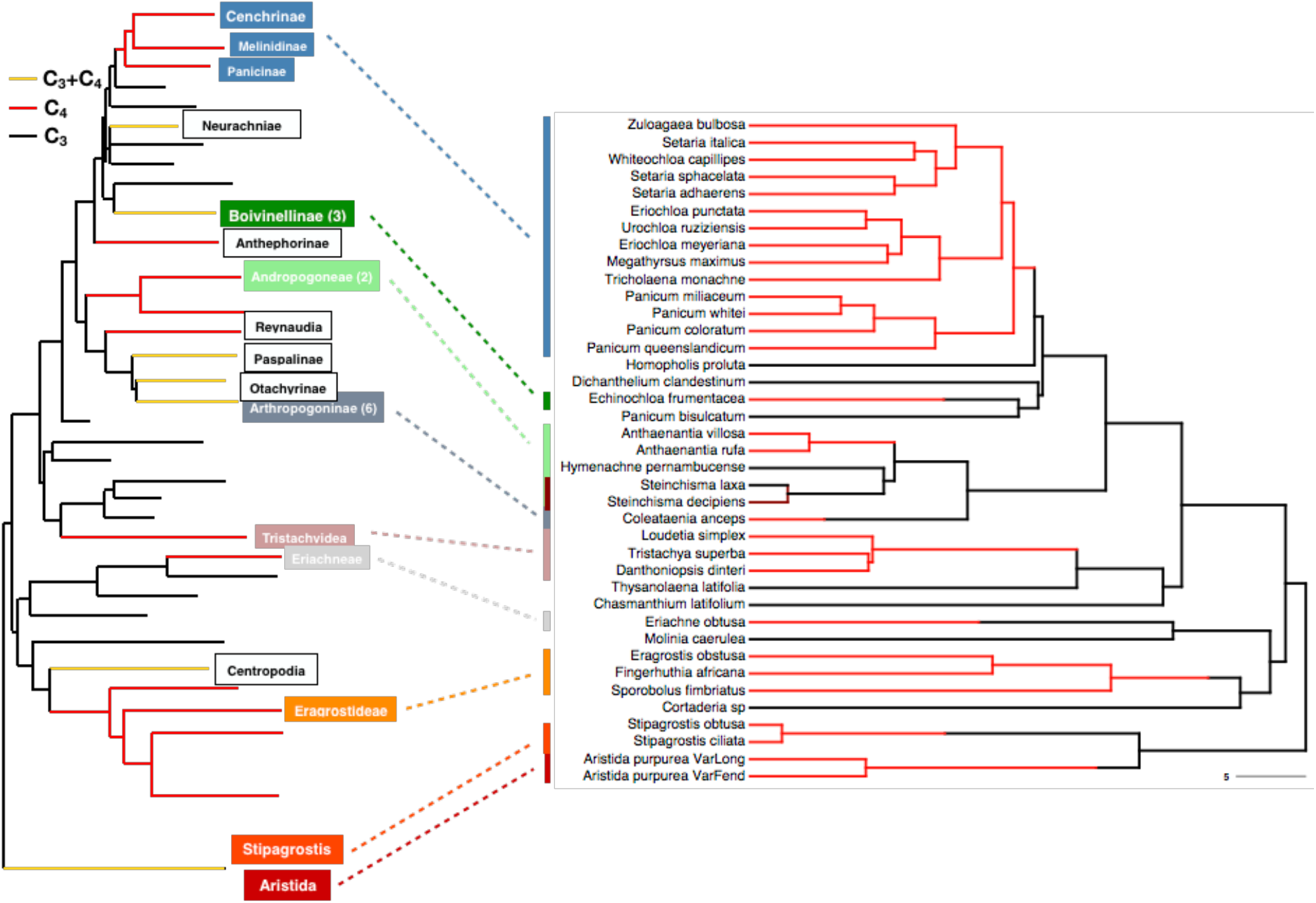
Phylogenetic sampling of the species for measuring physiological traits and the independent evolutionary lineages corresponding to grass lineages. The figure on the left was a grass phylogeny adapted from GPWGII (2012), on which the tags represent the recommended independent evolution of C_4_ for comparative studies in grasses (numbers represent there are multiple origins within a lineage). The figure on the right is the phylogeny for our species, extracted from a dated phylogeny ^[19]^ for species sampled in our experiments. We sampled nine independent evolution of C_4_ in total.

## Results

Within each phylogenetic cluster, there were no clear patterns between C_3_ and C_4_ hydraulic traits by conducting ANOVA tests only. C_4_ grasses had higher or equivalent K_leaf_, leaf capacitance leaf turgor loss point, A_max_ and lower or equivalent g_s_ than their closest C_3_ relatives (Fig. 2). The one C_3_-C_4_ intermediate species, *Steinchisma decipiens*, in our analysis had K_leaf_ similar or equivalent to C_4_, but leaf capacitance, leaf turgor loss point, g_s_ and A_max_ equivalent to C_3_ (Fig. 2). By analyzing our data in the context of the evolutionary models (Supplementary Table S1), however, we found clear C_3_-C_4_ differences in most measured traits. We first fitted evolutionary models of Brownian motion and Ornstein-Uhlenbeck processes to the hydraulic traits based on a reliable dated phylogenetic tree^[19]^. The best fitting evolutionary model to the data for K_leaf_, leaf turgor loss point, A_max_ and g_s_ was Ornstein-Uhlenbeck model, while the Brownian model is the best-fitting model for leaf capacitance, as determined by the AICc and Akaike weights and LRT test (Table 1, Supplementary Tables S2-S6). Higher K_leaf_, higher A_max_, lower leaf turgor loss point, and lower g_s_ are detected C_4_ species compared to C_3_ (LRT test, all *p*<0.01; all ΔAICc<-3). For leaf capacitance, there is no significant difference for C_3_ and C_4_ species.

**Fig. 2.**
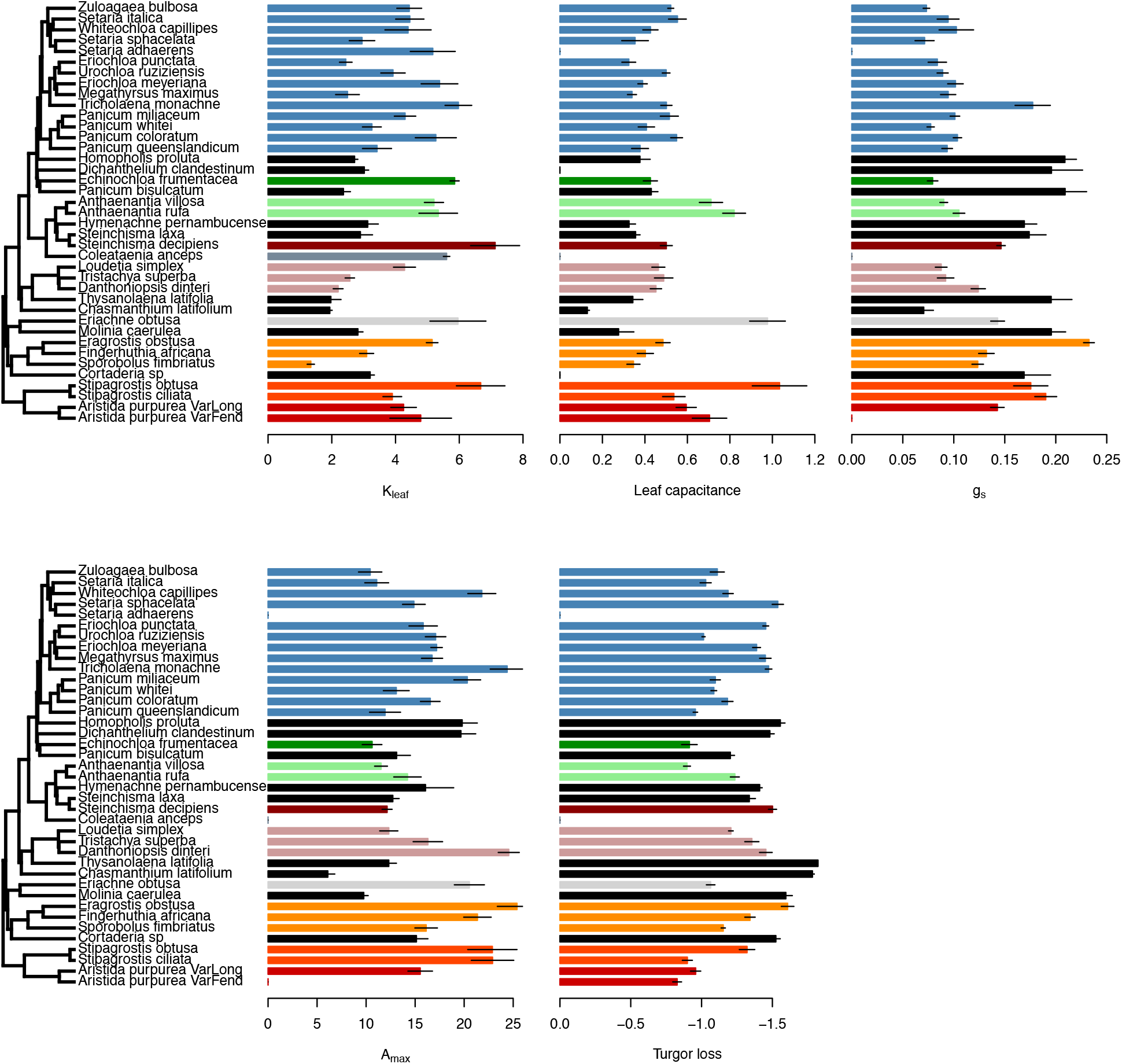
Hydraulic conductance (K_leaf_, mmol m^−2^ s^−1^ MPa^−1^), leaf capacitance (mmol m^−2^ MPa^−1^), maximal stomatal conductance (g_s_, mmol m^−2^ s^−1^), maximal assimilation rate (A_max,_ μmol m^−2^ s^−1^), and leaf turgor loss points (Turgor loss, -MPa) of closely related C_3_ and C_4_ species. Different colored clusters of bars show nine different origins of closely-related C_3_ and C_4_ species. C_3_ species are colored black. Error bars indicated standard errors.

**Table 1.**
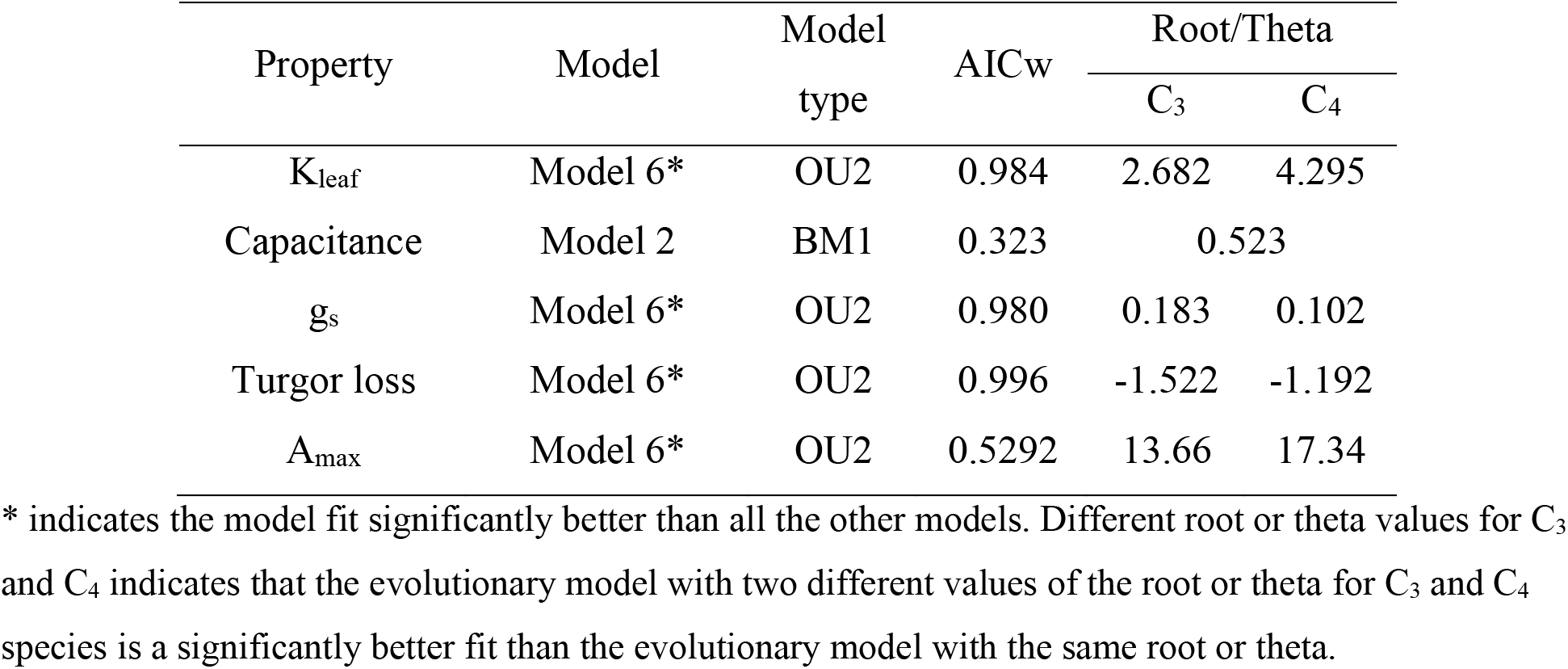
Phylogenetic results of the best-fitted models and their parameters for hydraulic conductance (K_leaf_), leaf capacitance (Capacitance), stomatal conductance (g_s_), and leaf turgor loss point (Turgor loss) (summarizing Table S2-S6; model description: Table S1).

We also looked for evolutionary trends in hydraulic traits after the evolution of a C_4_ system to probe for an extended ‘optimization’ phase of C_4_ evolution^[3, 20]^. Identifying directional trends in continuous character evolution is difficult without fossil taxa, and it is impossible to directly measure hydraulic traits for fossils; however, we can test for trends indirectly using extant species. For example, if reduction in K_leaf_ is selected for subsequent to C_4_ evolution we expect older C_4_ lineages to have lower K_leaf_ values than younger C_4_ lineages. We extracted the evolutionary age of C_4_ origin for each of our lineages from the dated phylogeny^[19]^. Regressions of evolutionary age versus hydraulic traits provide strong evidence for a long-term directional trend in hydraulic evolution following the origin of C_4_ photosynthesis (Fig. 3). K_leaf_, leaf turgor loss point and capacitance showed significant negative correlations with evolutionary age, while A_max_ had a significant positive correlation. In contrast, there was no significant relationship between g_s_ and evolutionary age. No evolutionary relationships were detected in C_3_ species, which indicated the correlations between evolutionary age and hydraulic traits were unique to C_4_ species. We also tested for an evolutionary trend by modelling hydraulic trait evolution using a phylogeny with branch lengths scaled to molecular substitutions/site, which provides an estimate of differences in evolutionary rates between lineages^[4]^. While the second approach requires many assumptions that are likely violated, the results also provide additional support to a directional trend in K_leaf_ and capacitance in C_4_ lineages: comparing 12 different types of models with or without evolutionary trends (supplementary Table S7), we found K_leaf_ and leaf capacitance were best fitted by the Brownian motion model with a significant negative trend for C_4_ (Supplementary Table S8, Table S9-13).

**Fig. 3.**
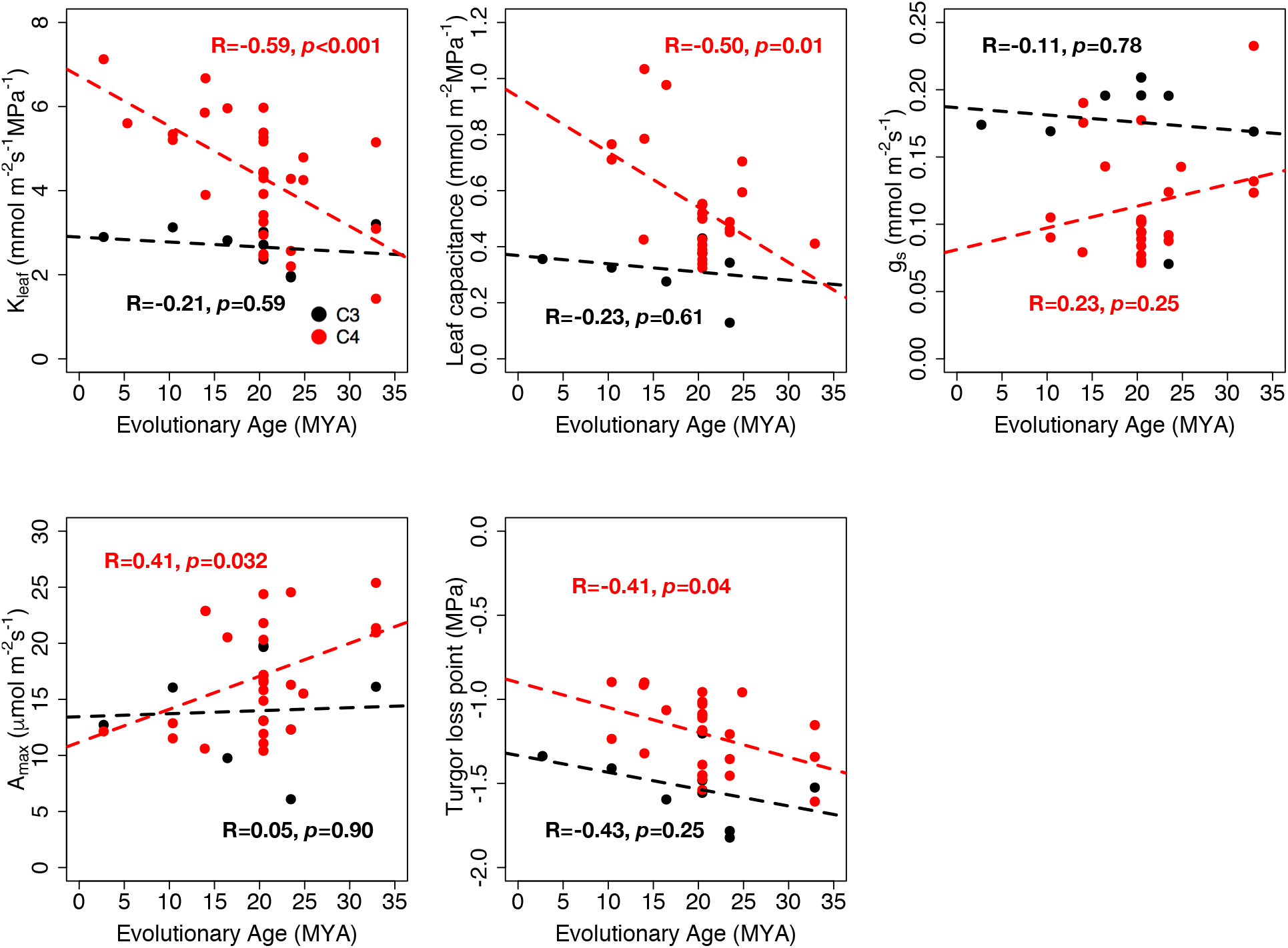
The regression for hydraulic conductance (K_leaf_), leaf capacitance, leaf turgor loss point, stomatal conductance (g_s_) and maximal assimilation rate (A_max_) vs. the evolutionary age for the nine origins of C_4_ to show the evolutionary trend within C_4_ and within their closely-related C_3_ species. The evolutionary age for each sampled origin is derived from the dated phylogeny^[19]^.

We next explored how A_max_ and hydraulic traits are correlated across the phylogeny, and whether this relationship is different for C_3_ and C_4_ lineages. The correlations between A_max_ and K_leaf_ were different between C_3_ and C_4_ (Fig. 4, Table 2, Table S13). A_max_ was significantly positively correlated with K_leaf_ for C_3_, but not for C_4_ (Fig. 4, Table 2, Table S13). A_max_ was weakly positively correlated with leaf capacitance and g_s_ and the correlations were not significantly different for C_3_ and C_4_ (Fig. 4, Table 2, Supplementary Table S21, S22). A_max_ was negatively, but not significantly related with leaf turgor loss point in C_3_ and C_4_ species (Supplementary Table S23).

**Fig. 4.**
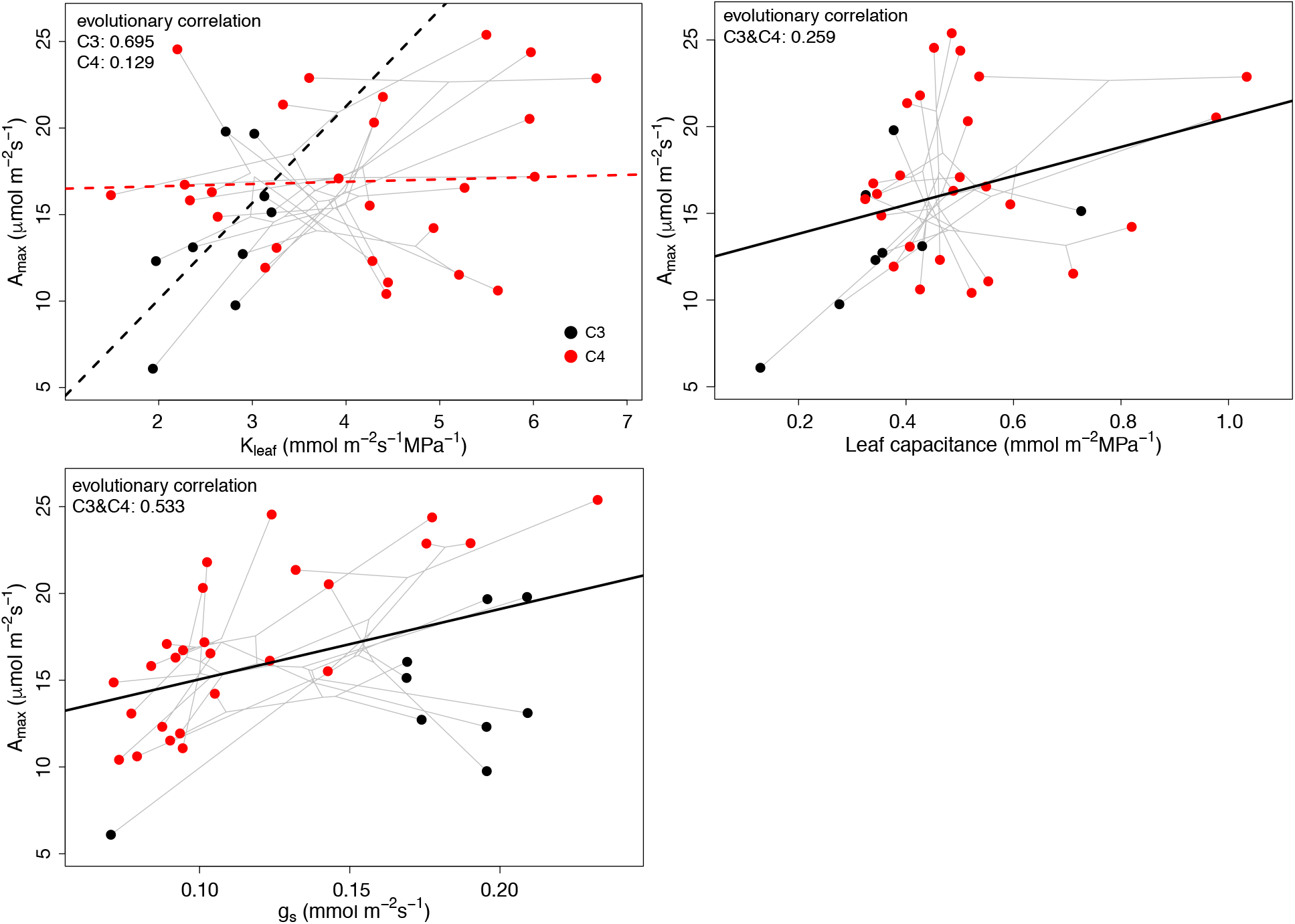
Phylogenetic correlation for C_3_ and C_4_ between A_max_ and other hydraulic traits (K_leaf_, leaf capacitance and g_s_). Different/same correlation values on the figure mean C_3_ and C_4_ have significantly different/same correlations. Detailed phylogenetic correlation models and analysis results are shown in Table 2. Dashed black line: C_3_; dashed red line: C_4_; solid black line: C_3_ and C_4_ have the same correlation; grey lines indicate the phylogeny.

**Table 2.**
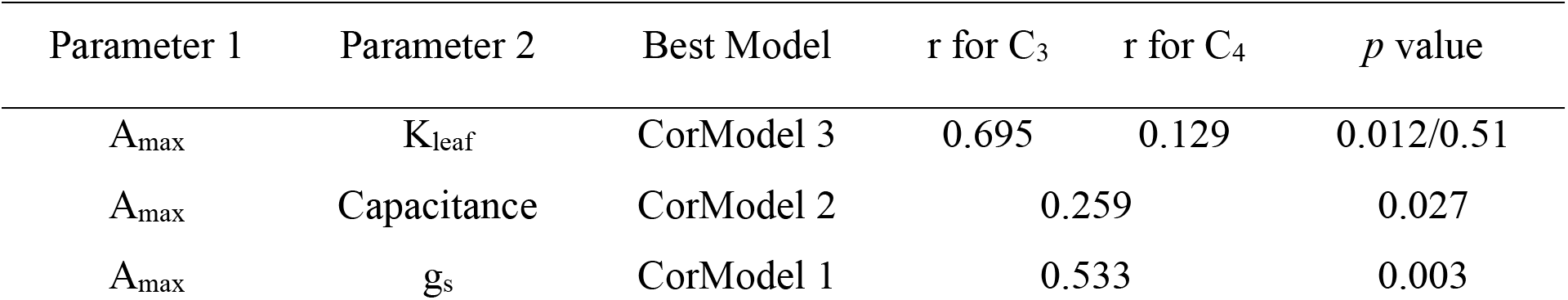
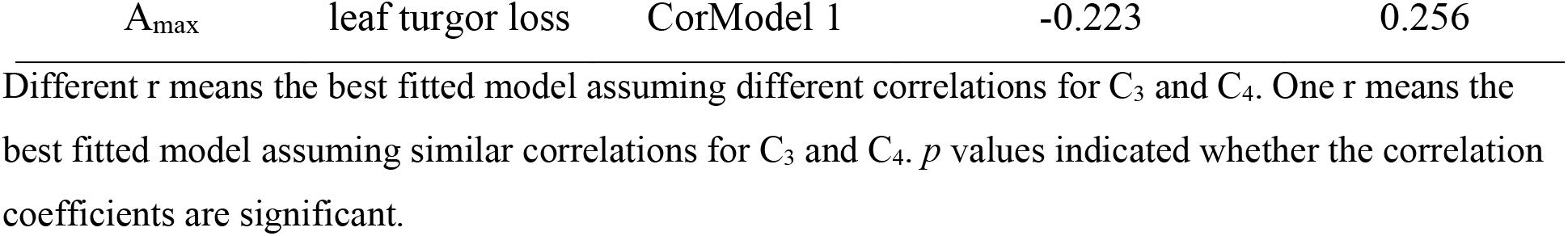
Phylogenetic Correlations between maximal assimilation rates (A_max_) and hydraulic traits for C_3_ and C_4_ species (summarizing Table S20-S23; model description: Table S19).

We used our mechanism-based physiological model^[32]^ to consider how the evolution of higher K_leaf_ would affect the optimal g_s_ and photosynthesis in C_3_ and C_4_ plants. An increase in K_leaf_ in the C_3_ ancestor selects for higher g_s_ and increases the steady-state leaf water potential to a limited extent (Fig. 5, S1). Changing K_leaf_ has a smaller effect on the photosynthesis rate of C_4_ than that of C_3_ (Fig. 6, Table S25), Decreasing K_leaf_ by half or doubling it changes the photosynthesis rate of a C_4_ plant by an average of −4.27% and 3.48%, respectively. In contrast, the same shifts in K_leaf_ has average effects of −10.07% and 9.14% on the assimilation rate of a C_3_ plant. The sensitivity of the assimilation rate to changes in K_leaf_ decreases with increasing CO_2_ concentration and increasing water-limitation for both C_3_ and C_4_ plants (Table S25). These differences in sensitivity to K_leaf_ were robust to differences in physiological properties between C_3_ and C_4_ (specifically, the temperature response properties and J_max_/V_cmax_ ratio; Table S25). The assimilation rate of C_4_ plants was still less sensitive to K_leaf_ than that of C_3_ species under different CO_2_ concentration and water-limited conditions (Table S25). The physiological modeling results indicates that C_4_ species maintain lower g_s_ and higher leaf water potential compared to closely related C_3_ species because the CCM reduces transpirational demand. The modeling effects of varying K_leaf_ on photosynthesis confirmed the diminished returns for high-efficiency water transport in C_4_ species mentioned above.

**Fig. 5.**
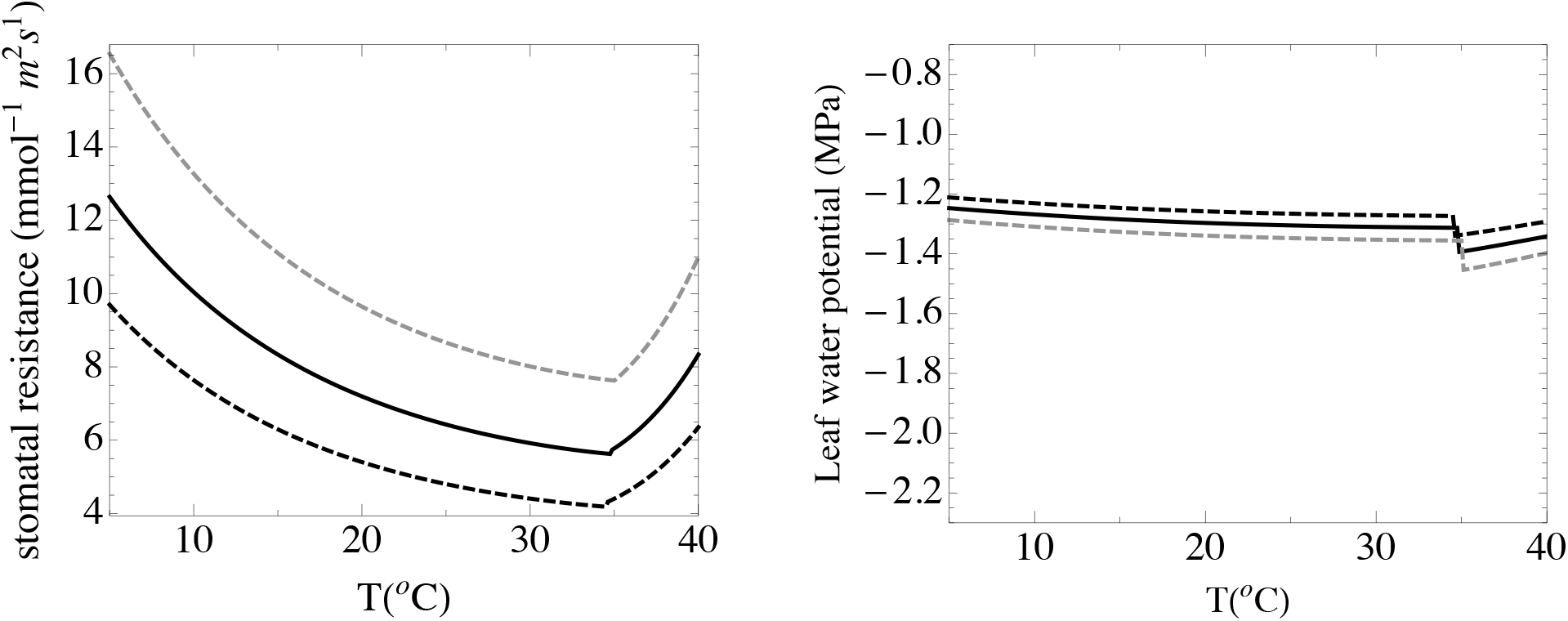
The effect of changing K_leaf_ on stomatal resistance (the inverse of g_s_) and leaf water potential under VPD=1.25 kPa, *ψ*_s_ =−1 MPa and CO_2_ concentration of 200 ppm for the C_3_ model. Solid black line: measured K_leaf_; dashed black line: K_leaf_ doubled; dashed grey line: K_leaf_ reduced by 50%.

**Fig. 6.**
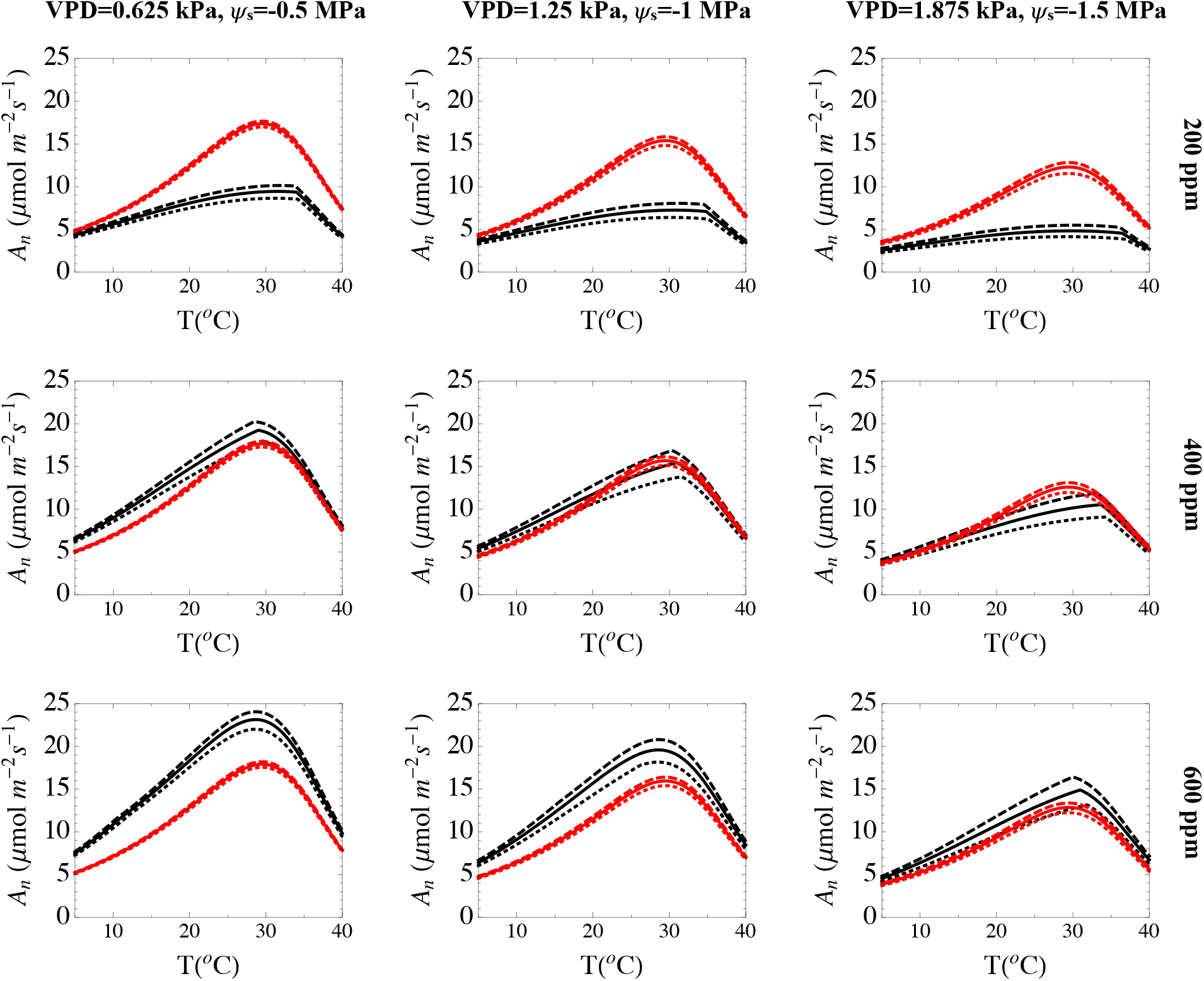
Modeling results of photosynthesis rates along with different CO_2_ concentration, different temperatures and different water limited conditions for C_3_ (black lines) and for C_4_ (red lines). Solid lines: modeling results for C_3_ and C_4_ with measured leaf hydraulic conductance; dashed lines: modeling results for C_3_ and C_4_ with twice of the regular leaf hydraulic conductance; dotted lines: modeling results for C_3_ and C_4_ with half of the regular leaf hydraulic conductance. C_3_ and C_4_ parameters are kept the same except for C_4_ has the carbon concentration mechanism.

To see if C_4_ subtypes varied in hydraulic traits and their evolutionary rates or variance, we also considered evolutionary models where we allowed each variable to have a subtype-specific value (Supplementary Table S1). We found no significant differences in K_leaf_, leaf capacitance, g_s_, leaf turgor loss point and A_max_ among C_4_ subtypes (all ΔAICc>0, ΔAICc obtained by AICc of subtype models minus AICc model not considering subtypes; Supplementary Tables S14-18). Although different decarboxylation enzymes are utilized by the three major subtypes (NADP-ME, NAD-ME and PCK), there does not seem to be an evolutionary effect on hydraulic traits. However, a previous study documenting PCK species from the Chloridoideae and Panicoideae lineages with lower leaf turgor loss point^[23]^. Such differences were not apparent when we compared C_4_ subtypes with multiple lineages. Our current representation of different subtypes is, however, somewhat limited. It would be advantageous to increase both lineage and species diversity and to balance subtypes within lineages to more deeply examine C_4_ subtypes.

## Discussion

The evolution of the C_4_ pathway in the grasses caused a series of shifts in hydraulic properties as compared to closely-related C_3_ grasses. The anatomical requirements of C_4_ initially increased K_leaf_ and leaf capacitance, as predicted by previous studies^[14,15,16]^; however, K_leaf_ and leaf capacitance appear to decline over evolutionary time, suggesting a long period of physiological optimization after the initial assembly of a new photosynthetic system. Previous examination of leaf hydraulic traits in grasses focused on investigating single species or were not developed within a phylogenetic framework when comparing multiple species^[21,22]^, and phylogenetic studies have assumed trait evolution as simple Brownian motion^[23,24]^. Hydraulic traits, however, may have evolved along different trajectories before and after the evolution of the C_4_ pathway and associated anatomical reorganization, resulting in more complicated evolutionary dynamics. Our evolutionary models indicated C_4_ grasses initially had higher K_leaf_, leaf capacitance, turgor loss point than corresponding C_3_, and a lower stomatal conductance (g_s_) than grasses consistent with previous studies^[25,26]^. Decreased vein distance and increased bundle sheath size are thought to be anatomical precursors to the evolution of C_4_^[27,28]^, and both are thought to increase K_leaf_ and/or leaf capacitance^[14,15]^. Therefore, the shifts of K_leaf_ and leaf capacitance likely occurred before, or at the initial formation of, the C_4_ CCM. After the full formation of C_4_, K_leaf_ and/or leaf capacitance started to decrease, which led to higher or equivalent K_leaf_ and leaf capacitance in the current C_3_ and C_4_ species (Fig. 2). Liu et al. (2019) found that K_leaf_ in C_4_ grasses overlapped with C_3_ values^[24]^. The positive correlation between A_max_ and the evolutionary age also supports an extended optimization phase for C_4_. Previous studies have indicated that species from the oldest C_4_ lineages (*Chloridoideae* and *Andropogoneae* for example) contain the most productive crops (Sage, 2016), while some recent C_4_ lineages are not more productive than C_3_ (Ripley et al., 2008; Lundgren et al., 2016). In contrast, the significant decrease of g_s_ and the increase of leaf turgor loss point occurred with the evolution of a fully operational C_4_ CCM, as suggested by our physiological models discussed below. Consistent with this prediction, in clades that possess a range of C_3_, C_3_-C_4_ intermediate and C_4_ physiologies, the increased water use efficiency, decreased g_s_, and a broadened ecological niche are observed only in plants with a full C_4_ CCM^[29,30]^.

The evolution of C_4_ significantly alters the widely-accepted A_max_-K_leaf_ relationships existing in vascular plants. A_max_ is limited by the efficient transport of water through leaves to replace water loss through open stomata, which is the likely cause of a positive correlation between K_leaf_ and A_max_ across and within plant taxa^[11,13,31]^. We found that A_max_ and K_leaf_ are positively correlated in our C_3_ species but not in C_4_ (Fig 4). Ocheltree et al. (2016)^[22]^ similarly found no relationship between K_leaf_ and A_max_ in a set of nine C_4_ species. We see possible explanations that are not necessarily mutually exclusive. First, the positive relationship of A_max_ and K_leaf_ is weakened under high K_leaf_, possibly due to diminished returns of further increasing the efficiency of water transport^[11, 31]^, a conclusion supported by our physiological modeling results below. As K_leaf_ tends to be lower in grasses than in other species, it is possible that the diminishing returns from increasing K_leaf_ manifest at lower values in grasses, and the initial high K_leaf_ resulting from C_4_ anatomy could be in the A_max_ “saturation” zone. Lastly, we see evidence here that the time-since-C_4_-evolution affects several hydraulic traits across and within lineages, and it could be that a walk towards A_max_–K_leaf_ optimality is slowly occurring within C_4_ grass lineages in relatively newfound ecological niches. However, the similar correlations of g_s_ vs. A_max_ in C_3_ and C_4_ and lack of evolutionary trend in g_s_ indicated the evolutionary processes of g_s_ might be already near the optimal condition or stabilized quickly. Other hydraulic traits of leaf capacitance and leaf turgor loss point do not seem to contribute to the A_max_ directly because of weak correlations.

We identified the mode and direction of evolution for hydraulic traits in C_3_ and C_4_ lineages and found evidence that different traits followed different evolutionary processes. Hydraulic conductance and leaf capacitance could therefore evolve with directions in a step-wise fashion due to anatomical constraints, but g_s_ and leaf turgor loss point might have a more quick process of readjustments, which allows them to stabilize soon. This suggests that there could be greater diversification of K_leaf_ and leaf capacitance in the existing C_4_ species and maybe in the future. Also, these rearrangements of hydraulic properties interacted with each other throughout the evolutionary trajectory. For example, increased K_leaf_ and leaf capacitance would lead to an increased water transport efficiency, which enabled greater g_s_ of the C_4_ ancestor (either a C_3_ grass or a C_3_-C_4_ intermediate), but the formation of the full C_4_ CCM enables a decrease of g_s_. Therefore, observed g_s_ in C_4_ grasses reflects a balance of these two contrasting physiologies playing out in a given ecological and phenological background, which may explain why although C_4_ g_s_ was lower than the C_3_, the difference was not large. This line of reasoning might also explain the inconsistent observations of g_s_ comparisons between C_3_ and C_4_. Most previous studies found that C_4_ grasses had lower g_s_ than C_3_ grasses in both closely related and unrelated species^[25,33]^, yet Taylor et al. (2014) found that C_4_ grasses maintained a higher or equivalent g_s_ to closely-related C_3_ grasses^[34]^. Likewise, artificial selection or genetic engineering might have more success in adjusting these hydraulic traits in advance. Consciously selecting or manipulating narrower xylem, decreasing the expression of aquaporins, or other mechanisms of decreasing leaf conductance while maintain high bundle sheath to mesophyll ratio, together with CCM may increase the water use efficiency of C_4_ species further. Our phylogenetic analyses can thus inform both the evolutionary history of C_4_ plants and future efforts to modify C_4_ crops.

By capitalizing on the multiple origins of C_4_ photosynthesis in grasses, we have shown that the vascular organization that is a hallmark of C_4_ plants also impacts leaf hydraulics, and disrupts the established link between hydraulic and photosynthetic capacity demonstrated in C_3_ plants. C_4_ grasses are “overplumbed” relative to their C_3_ counterparts, suggesting that the costs associated with the production of an extensive leaf vasculature require re-evaluation in plants with C_4_ photosynthetic systems. The gradual decline in K_leaf_ in C_4_ lineages over millions of years also requires an explanation. The C_4_-K_leaf_ conundrum provides an opportunity to examine what we mean by “evolutionary constraint” and highlights the very dynamic nature of evolutionary trade-offs and functional optimization. First, we assume that the costs of building and maintaining a high K_leaf_ are still significant in C_4_ plants^[12,35,36,37,38]^. The most efficient way to reduce K_leaf_ costs would be to reduce venation density, as veins come with high construction costs^[12,17]^, and also reduce the leaf area that is available for carbon fixation. Yet the anatomical requirements of the C_4_ system preclude this option: reducing vein density would result in a highly inefficient C_4_ system^[15]^, which would negatively impact the plant’s carbon budget, presumably to a much greater extent than the cost of an overbuilt venation system. As vein construction is a primary contribution to the cost of a high K_leaf_, and high vein densities are now linked to a new function (C_4_ carbon fixation), the cost-benefit calculations in optimizing K_leaf_ have shifted, and the tradeoff is in favor of overplumbing in order to maintain a highly efficient new carbon fixation system. In evolutionary vocabulary, what emerges is a new constraint – and in this example, it is clear that the emergence of a new constraint to organismal evolution is simply due to a shift in the tradeoffs associated with characters that influence multiple aspects of organismal function. In other words, we assume a low vein density is a phenotype that is still developmentally achievable for C_4_ grasses; what has prevented its emergence is the shift in functional costs associated with reduced vein densities.

And yet, we documented a gradual reduction in K_leaf_ over time, which we presume was accomplished via changes in other factors that influence leaf hydraulic capacity– perhaps by changing xylem conduit diameters, shifts in extra-xylary mesophyll conductance, decreased expression of aquaporins, and reorganization of internal air spaces^[6,12,37,39,40]^. It is possible that these changes resulted from a continued and direct selection pressure to reduce investment in an underutilized hydraulic system. An alternative explanation is that all of the traits that influence K_leaf_ also play important roles in other aspects of leaf function – and the emergent of a new constraint (a high vein density to maintain C_4_ function) has ***released*** still other constraints on other traits so that they may be optimized for their other functions. A striking pattern in our data is that older C_4_ lineages have achieved both lower K_leaf_ and higher A_max_ – suggesting that they are continuing to optimize their photosynthetic capacity, long after the initial origin of C_4_. We suspect that the slow evolutionary decline in K_leaf_ is due in large part to the optimization of traits to increase A_max_ at the expense of K_leaf_, which is possible only because hydraulic capacity was already “buffered” by the vein density requirements of C_4_ – allowing for continued reductions of K_leaf_ at no functional cost. Increased suberization of bundle sheath cells is one example of a potential release of constraint^[22]^: it allows C_4_ plants to gain higher A_max_ through reducing bundle sheath leakiness, but it likely simultaneously reduces water flow from veins out into the mesophyll. Since C_4_ plants are already operating in hydraulic excess, bundle sheath suberization may be optimized for C_4_ function without any negative repercussions for plant water relations. This hypothesis could also explain the opposing trends in A_max_ and K_leaf_ when viewed as a function of evolutionary age. The examination of C_4_ evolution in grasses provides an exciting system to study the evolutionary dynamics of constraints highlighted by the interplay between photosynthesis and plant hydraulics.

## Methods

### Plant material

We collected seeds of 39 closely related C_3_ (9 species), C_4_ species (29 species), representing three C_4_ subtypes, nine C_4_ origins, and one C_3_-C_4_ intermediate species. The selected C_3_ and C_4_ species fall into nine identified C_4_ lineages belong to the 11 recommended grass lineages for C_3_ and C_4_ study (11 out of the total 24 grass lineages have clear C_3_ sister species and are recommended for comparative studies in GPWGII, 2012^[4]^): *Aristida*, *Stipagrostis*, *Chloridoideae* (*Eragrostideae*), *Eriachne*, *Tristachyideae*, Arthropogoninae, *Otachyrinae (Anthaenantia)*, *Panicinae*, *Melinidinae*, and *Cenchrinae* (Fig. 1). In 2015, seeds were surface sterilized before germination and the seedlings were transferred to 6 inch pots with the soil of Fafard #52 (Sungro, Ajawam, MA). Six replicates of each species were randomized in the greenhouse of the University of Pennsylvania supplemented with artificial lighting. The plants were watered twice daily. Daytime/night temperature was controlled at 23.9-29.4/18.3-23.8°C; relative humidity was around 50-70%. Plants were fertilized once per week with 300 ppm Nitrogen solution (Jacks Fertilizer; JR Peters, Allentown, PA) and 0.5 tsp of 18-6-8 slow release Nutricote Total (Arysta LifeScience America Inc, NY) per pot was applied when plants were potted into 6 inch pots. To maintain optimal plant growth a 15-5-15 cal-mg fertilizer was used every third week. All measurements were performed on the most-recent fully expanded leaves.

### Hydraulic traits

Leaf hydraulic conductance (K_leaf_) was measured using the evaporative flux method^[41]^, with some adjustments to maintain stability of the evaporative environment to which the leaf was exposed (Supplementary Methods). The evening before measurements, potted plants were brought to the laboratory, watered, and then covered by black plastic bags filled with wet paper towels to rehydrate overnight. For the leaf gasket, a 1 cm diameter, ~ 1 cm long solid silicone rubber cylinder was cut nearly in two, leaving a hinge on one end. The cylinder was placed around the leaf blade near the ligule and glued shut with superglue^[42]^. The leaf was cut from the plant with a razor blade while submerged in a 15 mmol L^−1^ KCl solution; the rubber gasket was then attached to tubing filled with the same KCl solution. The other end of the tubing was inside a graduated cylinder that sat on a digital balance (Mettler-Toledo). The leaf was then placed inside a custom, environmentally controlled cuvette that allowed for the measurement of entire grass blades. Throughout measurements, cuvette temperature was controlled at 25 °C and the humidity was 55-65% (VPD range of 1.1-1.4 kPa) across measurements, but remained constant during a particular measurement. Photosynthetically active radiation in the system is 1000 μmol m^−2^ s^−1^. Flow from the balance was monitored for 45 m to 1h until the flow rates reach steady state. After the measurements, the leaf was detached and was put into a plastic bag to equilibrate for 20 minutes to measure the leaf water potential (Model 1000, PMS Instrument, USA). K_leaf_ values were further standardized to 25°C and leaf area to make the K_leaf_ comparable among studies and across species. Data indicating a sudden change of flow and whose leaf water potential was an obvious outlier were deleted.

We measured pressure-volume (PV) curves for six leaves per species using the bench-drying method^[43,44]^. A leaf was cut directly from the same plants rehydrated in the lab (as described above) using a razor blade and leaf water potential was measured immediately. Then, the leaf weight was recorded. The leaf was initially allowed to dry on the bench for 2-minute intervals and put into a ziplock bag and under darkness for 10-minute equilibration before measuring the leaf water potential and leaf weight again. Then, the waiting intervals could be adjusted based on the decrease of the leaf water potential (from 2 minutes-1h). Ideally, a decreasing gradient of −0.2MPa for leaf water potential was obtained for the curves, until the leaf weight reached a steady state. At the end of the experiment, leaves were dried in the oven at 70°C for 48h to obtain the dry weight. The PV curves were used in curve fitting to obtain leaf capacitance, and leaf turgor loss point using an excel program from Sack and Pasquet-Kok (2010)^[44]^.

Maximal assimilation rate (A_max_) and stomatal conductance (g_s_) were measured under saturated light intensity. A_max_ and g_s_ were obtained using a standard 2 × 3 cm^2^ leaf chamber with a red/blue LED light source of LI-6400XT (LI-COR Inc., Lincoln, NE, USA). Light curves were measured with light intensities of 2000, 1500, 1200, 1000, 800, 500, 300, 200, 150, 100, 75, 50, 20, 0 μmol m^−2^ s^−1^ under CO_2_ of 400 ppm. Then, A_max_ was estimated from the light curve^[45,46]^. All the measurements were made under the temperature of 25°C and the leaf temperature to air vapor pressure deficit was controlled around 2kPa. g_s_ at the saturated light intensity of 2000 μmol m^−2^ s^−1^ was recorded for each plant. The cuvette opening was covered by Fun-Tak to avoid and correct for the leakiness.

### Phylogenetic analysis

#### Phylogenetic analysis for C_3_ and C_4_

We pruned the dated phylogeny from a published grass phylogeny to include only the species in our physiological experiments^[19]^(Fig. 1). Using the dated phylogeny, for each of the hydraulic traits, we fitted evolutionary models to test which evolutionary model best explains observed distribution of traits along the phylogeny and how these models differ between C_3_ and C_4_ (Table S1). We fitted evolutionary models belonging Brownian Motion model and Ornstein-Uhlenbeck Model using the package “mvMORPH” in R^[47]^. To determine the best fitted evolutionary model, we compared two criteria, the small-sample-size corrected version of Akaike information criterion (AICc, the lower AICc, the better fit) and Akaike weights (AICw, the higher AICw, the better fit)^[48,49,50]^. The evolutionary models have nested variants (Models 1-4; Models 5-6), varying in whether C_3_ and C_4_ species had the same or different fluctuation rates, root states for Brownian motion model and optima for Ornstein-Uhlenbeck model. We used likelihood-ratio test (LRT) to verify whether a specific model variant performs significantly better. The AICc, AICw and LRT allowed us to test evolutionary hypotheses, for instance, if the model in which C_3_ and C_4_ have different root states fit significantly better than model in which C_3_ and C_4_ have the same root states, it means there is a shift of physiological trait along with the formation of C_4_. To examine the further evolution of hydraulic traits after a full C_4_ evolved, we extracted the evolutionary ages for each represented C_4_ origin from the dated phylogenetic trees. Then, we regressed the hydraulic traits with evolutionary age. A significant negative correlation between evolutionary age and hydraulic trait will indicate a further decreasing evolutionary direction after C_4_ evolved. We also performed an additional analysis to test the original states and further direction together. We extracted molecular phylogeny for all the species from Edwards, GPWG II (2012)^[4]^. Except for the six evolutionary models mentioned above, the molecular phylogeny allows us to fit for additional six Brownian motion models with trend (Supplementary Table S7). Likewise, if Brownian motion model with trend fits the phylogenetic patterns better than Brownian motion model without trend it means there is an evolutionary trend, and a significant LRT test for a two-trend model suggests that C_3_ and C_4_ lineages differ in the speed or direction of hydraulic evolution. We also mapped the traits on the phylogeny for potential further references (Fig. S2-S5).

To further test whether there are significant differences among C_4_ subtypes, evolutionary models with subtypes (Table S1) were used to fit the data. We again used AICc, AICw and LRT methods to find the best model variants: whether there are significant differences for hydraulic shifts and evolutionary trends among three different subtypes. For the leaf capacitance analysis, *Dichanthelium clandestinum* is deleted as it is an obvious outlier.

#### Phylogenetic analysis for correlations among traits

Multivariate analysis in “mvMORPH” was used to estimate the correlations between A_max_ and each of the hydraulic traits and to test the hypotheses that whether such correlations are different between C_3_ and C_4_. The process of brownian motion with different root for C_3_ and C_4_ was used for K_leaf_, g_s_ and leaf turgor loss and brownian motion with the same root was used for leaf capacitance. Since the Ornstein-Uhlenbeck process is difficult to take the root state difference into consideration, here we used Brownian motion assumptions as approximation for leaf turgor loss. Seven different correlation models are fitted (Table S19). We used LRT for the seven correlation models to test whether the correlation of the two traits is significantly different from 0 and whether the correlation of two traits is significantly different between C_3_ and C_4_. Such correlation analysis is similar to PGLS considering C_3_ and C_4_, but with more varieties on the setting of variance and covariance matrix.

### Physiological Modeling

Furthermore, we used physiological models that couples the photosynthesis systems and hydraulic systems to predict the effect of changing K_leaf_ on assimilation rate^[32]^. The change of K_leaf_ was assumed to change the plant hydraulic conductance (K_plant_) proportionally in the modeling process. We double or reduce by half K_leaf_ relative to the original value to predict the effects on assimilation rates for C_3_ and C_4_ pathways. We assumed C_4_ had the same photosynthetic properties with C_3_ species (e.g., Rubisco affinity and specificity, Supplementary Table S24) other than the carbon concentration mechanism, which mimics the initial evolution of C_4_ and the closely-related C_3_-C_4_ system. We also model the additional scenarios in which C_4_ had different photosynthetic properties to support the above condition further (Supplementary Table S25).

## Acknowledgements

HZ and this research is supported by the NOAA Climate and Global Change Postdoctoral Fellowship Program, administered by UCAR’s Cooperative Programs for the Advancement of Earth System Science (CPAESS) under award #NA18NWS4620043B and is also supported by the Dissertation Completion Fellowship provided by the Graduate Division of School of Arts and Sciences, University of Pennsylvania. BH is supported by NSF-IOS award 1856587.

## Data availability

The data that support the findings of this study are available from the corresponding author upon request.

## Code availability

All source code is available upon request.

